# Long-term dysregulation of histone acetylation, BDNF expression, and the hippocampal transcriptome in adult rats exhibiting a depressive-like phenotype following repeated chlorpyrifos exposure

**DOI:** 10.64898/2026.11.02.763669

**Authors:** Ana Ribeiro-Davis, Elisa Hawkins, Fay M Jahr, Mikhail G. Dozmorov, Joseph L. McClay, Laxmikant S. Deshpande

## Abstract

Epidemiological evidence indicates that occupational exposures to the organophosphate pesticide chlorpyrifos (CPF) in adults are associated with an increased risk for depression. In this study, we use a rat model of occupational-like CPF exposure associated with a chronic depressive phenotype to investigate underlying molecular mechanisms. Adult rats received CPF (5 mg/kg/day) for 21 days. When assessed three months later, blood and brain cholinesterase activity had returned to control levels despite the presence of depressive signs. Hippocampal tissue was collected at this endpoint for molecular analyses. Western blotting analysis showed significant upregulation of histone deacetylase protein levels of HDAC3 and HDAC5, whereas HDAC1 and HDAC2 were unchanged. HDAC5 regulates histone acetylation at the brain-derived neurotrophic factor (*Bdnf*) gene. We used chromatin immunoprecipitation to assay histone 3 lysine 9 acetylation (H3K9ac) occupancy at the *Bdnf* promoter IV. H3K9ac levels at the *Bdnf* promoter IV were significantly reduced in CPF rats, accompanied by a concomitant reduction in BDNF protein expression. Exploratory RNA-sequencing identified 100 significantly differentially expressed genes in CPF-exposed rats at a false discovery rate < 5%, including significant downregulation in genes such as insulin-like growth factor 2 (*Igf2*), fibroblast growth factor 14 (*Fgf14*), and slit guidance ligand 2 (*Slit2*) involved in neuronal growth, synaptic function, and axon guidance. Gene Ontology analysis revealed enrichment in pathways related to neurogenesis, neuronal projections, and myelination. Overall, this study indicates persistent epigenetic and transcriptional dysregulation associated with reduced hippocampal BDNF and alterations in neuronal plasticity-related pathways linked to a long-term depressive-like phenotype following CPF exposure.

## Introduction

Chlorpyrifos (CPF) is one of the most widely used organophosphate (OP) pesticides, although its use varies geographically. While the European Union and Canada have implemented bans on CPF (Sami et al. 2025), its use remains substantial in China and India (Foong et al. 2020). In the United States, CPF regulation has fluctuated (Sami et al. 2025) and it remains registered for use on 11 high-value fruit and grain crops, along with several non-food applications, including turf management, commercial ornamentals and forestry, pest control, and industrial wood treatment (US EPA 2026). Consequently, workers and residents near these industries are exposed to CPF (Mackenzie Ross et al. 2010; Dereumeaux et al. 2020; Vaezafshar et al. 2024).

Concerns over the health impacts of CPF have grown in recent decades. Epidemiological studies, including the Agricultural Health Study, have identified associations between sustained low-level OP exposure and depression, anxiety, and suicidal ideation in agricultural workers, farm residents, and related rural populations (Stallones and Beseler 2002; Beseler et al. 2008; Beard et al. 2014; Harrison and Mackenzie Ross 2016; Joo and Roh 2016). While a correlation between gestational or early-life OP exposure and later-life neuropsychiatric problems was already established (Sagiv et al. 2021; Sagiv et al. 2023), these studies indicated that adult OP exposure could also give rise to neuropsychiatric problems. In this regard, Gulf War veterans represent a unique population who were chronically exposed to OP pesticides, including CPF and malathion, during their deployment. Upon return, many developed Gulf War Illness, a chronic multi-symptom condition that includes mood and memory disturbances (Haley et al. 1997; Kang et al. 2009). An unresolved problem with OP exposures is why the neurobehavioral deficits persist long after the exposures have subsided (Harrison and Mackenzie Ross 2016).

All OPs, including CPF, inhibit acetylcholinesterase (AChE) (Pope et al. 2005), resulting in synaptic acetylcholine accumulation. Depending on the OP compound, dose-dependent cholinergic toxicity can range from mild symptoms like miosis, salivation, and headaches to seizures and even death (Worek et al. 2016). Occupational or environmental exposure may cause partial AChE inhibition without acute hypercholinergic signs, yet low-level OP exposures are associated with chronic neuropsychiatric problems, including depression and cognitive dysfunction (Davies et al. 1999; Wesseling et al. 2002; Harrison and Mackenzie Ross 2016).

Epigenetic mechanisms have emerged as a plausible explanation for how transient environmental exposures can result in sustained biological and behavioral alterations. DNA methylation and histone post-translational modifications are among the epigenetic mechanisms that switch genes “on and off” in a cell (Sweatt 2013). Many factors affect epigenetic states, such as our environment and experiences (Collotta et al. 2013). Toxin exposures, including chronic low-level OP exposure, have been linked to epigenetic changes in human studies (Paul et al. 2018; Trivedi et al. 2019; Chiu et al. 2021) and in laboratory models (Kim et al. 2016; Ventura et al. 2019; Barzi et al. 2020; Sai et al. 2020; Ribeiro et al. 2021). Thus, given the relationship between OP exposures and depression, OP-induced epigenetic changes may therefore contribute to adverse CNS effects after exposure has subsided.

Prior research has reported delayed cognitive and synaptic effects of CPF exposure (Cañadas et al. 2005; Speed et al. 2012), whereas studies on molecular effects of repeated CPF exposure have largely focused on alterations during or shortly after CPF dosing (Yan et al. 2012; Lee et al. 2016; Voorhees et al. 2019). However, persistent molecular alterations associated with long-term depressive outcomes following occupational-like CPF exposure remain poorly understood. We recently reported that adult male rats injected with CPF (1, 3, 5, or 10 mg/kg/d, s.c.) for 21 days exhibited dose-dependent increases in signs of depression, including anhedonia, anxiety, and despair, when assessed after a 12-week washout period (Ribeiro et al. 2022). In this study, we investigate the molecular mechanisms underlying occupational-like CPF exposures and the elevated risk of chronic behavioral deficits. We focus on the 5 mg/kg/day for 21 days CPF dosing regimen because it produced acute blood cholinesterase inhibition consistent with those seen in human occupational exposures, based on PK/PD modeling (Smith et al. 2009; Ellison et al. 2011; Lein et al. 2012), along with substantial cholinesterase inhibition in the hippocampus and cortex (Ribeiro et al. 2022). Importantly, cholinesterase activity in both blood and brain had recovered by three months after exposure, even though the animals exhibited a robust depressive-like phenotype across multiple behavioral tests (Ribeiro et al. 2022). Thus, this model is well-suited to investigate persistent molecular alterations associated with long-term neurobehavioral consequences of CPF exposure after AChE inhibition has resolved.

## Materials and Methods

### Animals

All animal use procedures were in strict accordance with the National Institutes of Health (NIH) Guide for the Care and Use of Laboratory Animals. The experimental protocol was approved by Virginia Commonwealth University’s (VCU) Institutional Animal Care and Use Committee (IACUC). Male Sprague-Dawley rats (Inotiv, Indianapolis, IN) were ordered at 8 weeks of age and allowed to acclimate to the housing environment for 1 week. Animals were housed one per cage at 20-22 ^°^C with a 12-hour light-dark cycle (lights on 0600-01800 h), provided with appropriate environmental enrichment (nylon bone), and given free access to food and water.

### CPF exposure

Rats were 9 weeks old at the initiation of dosing, and initial body weights ranged from 240 to 290 g. Animals were monitored daily for weight or discomfort throughout the study. Rats were randomized into two groups of 10 rats. CPF (CAS no. 2921-88-2) was obtained from Chem Service, Inc. (Chester, PA) as a colorless crystalline solid (purity > 99.4%). CPF formulations in peanut oil (Millipore Sigma) were prepared twice weekly. CPF formulation (5 mg/kg; CPF-5) or an equal volume of the vehicle (peanut oil) was administered via subcutaneous (s.c.) injection for 21 consecutive days (study days 1-21), approximately 24 h apart. The injections were given along the back, and the administration sites were staggered to prevent local inflammation or buildup of subcutaneous oil depots (Levoe et al. 2014). These exposures were carried out in their home cage. Rats underwent behavioral testing at the beginning of week 12. As reported in Ribeiro et al. (2022), CPF-5 rats exhibited significant signs of anhedonia, anxiety, and despair, suggesting the presence of a depressive phenotype. At the end of the behavioral studies, rats were euthanized, brains were removed, the hippocampus was dissected, flash-frozen, and stored at -80 ^°^C. The experimental timeline is shown in **Fig. 1**.

**Figure 1.**
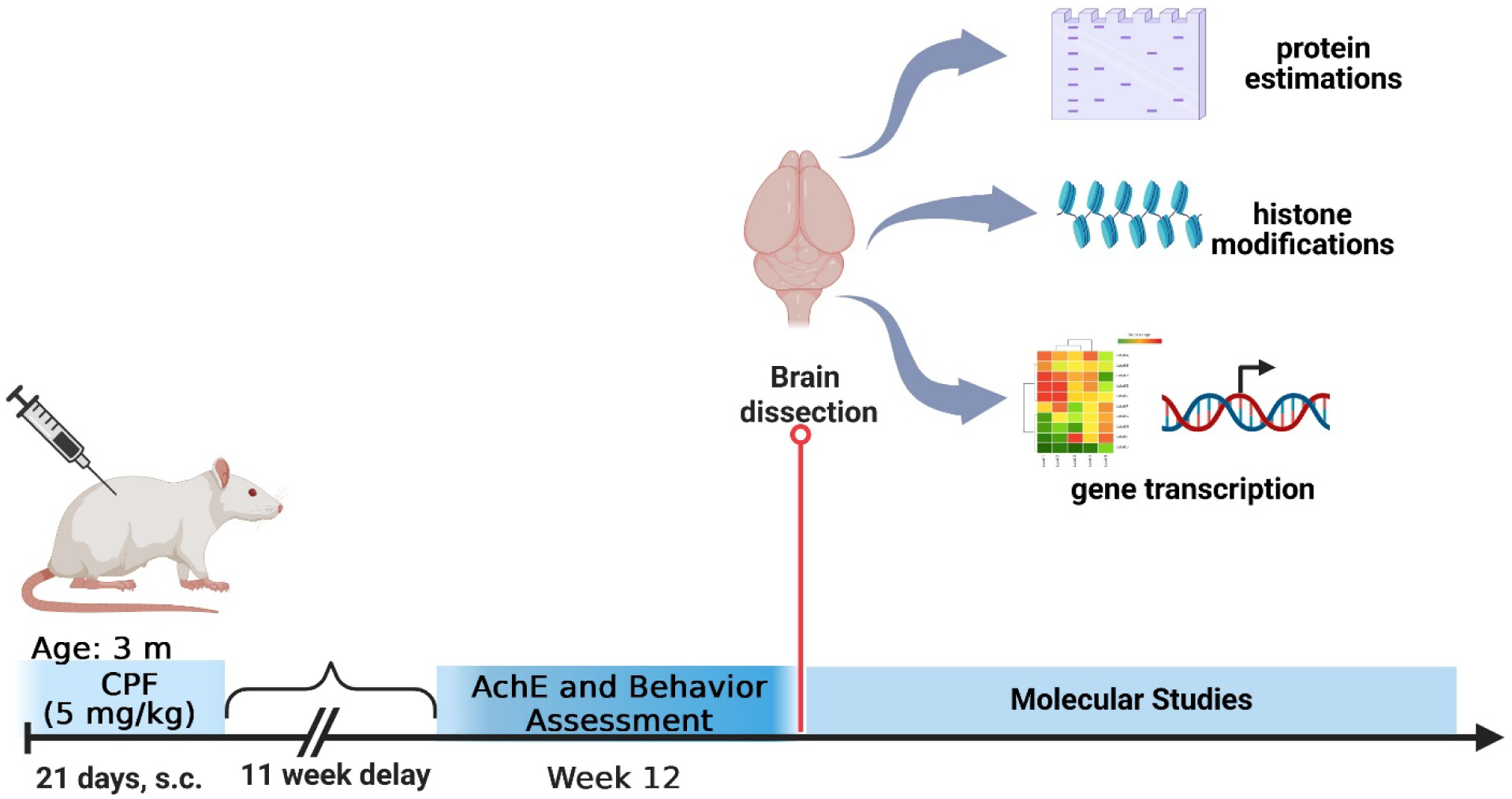
Experimental design for assessment of long-term molecular effects of CPF exposure. Male Sprague-Dawley rats (3 m old) received either CPF (5 mg/kg/d, s.c., 21 days) or vehicle (VEH; peanut oil). Signs of depression (anhedonia, anxiety, despair) were assessed at 12 weeks after exposure (Ribeiro et al. 2022). CPF and VEH rats were subsequently euthanized, and hippocampal tissue was micro-dissected for molecular studies.

### Western blots

Antibody-specific Western blotting was conducted using procedures established in our lab and validated for OP exposures (Ribeiro et al. 2020; Ribeiro et al. 2021). The blots were carefully cut between HDAC, BDNF, and Glyceraldehyde-3-Phosphate Dehydrogenase (GAPDH) bands based on visual identification using a Ponceau S temporary stain solution (P7170, MilliporeSigma). The HDAC sections were subsequently subjected to primary antibodies with the following dilutions: HDAC1 at 1:1500 (PA1-860, ThermoFisher), HDAC2 at 1:2000 (ab32117, Abcam), HDAC3 at 1:5000 (ab32369, Abcam), and HDAC5 at 1:1000 (sc-133106, Santa Cruz). In the case of the BDNF section, a primary BDNF antibody at 1:1000 (ab108319, Abcam) was employed. Meanwhile, the GAPDH section was probed with a 1:20000 dilution of the primary antibody (MA5-15738, Thermo Scientific) and incubated overnight at 4 °C. Subsequently, three 5-minute washes with TBST were performed. The blots were then exposed to secondary antibodies for detection, specifically horse-radish peroxidase (HRP)-coupled goat anti-rabbit IgG (1:10000, sc-2030, Santa Cruz) or goat anti-mouse IgG (1:10000, sc-2031, Santa Cruz) for the HDACs. The BDNF blot used a horse-radish peroxidase (HRP)-coupled rabbit IgG secondary antibody (1:10000, 7074S, Cell Signaling), while the GAPDH blot utilized HRP-coupled goat anti-mouse IgG secondary antibody (1:20000, sc-2031, Santa Cruz) for 1 hour at room temperature. Following this, the blots were washed five times with TBST for 5 minutes each. To initiate chemiluminescence, 2µL of Clarity^TM^ Western Enhanced chemiluminescence (ECL) Substrate (170-5060, Bio-Rad, Hercules, CA) was applied, and imaging occurred 5 minutes later. The individual blot sections were then realigned and imaged using the default auto-optimal exposure or manual exposure settings for up to 5 minutes, capturing an image every 10 seconds using a ChemiDoc Touch (Bio-Rad). Image analysis was conducted using ImageLab (Bio-Rad). The intensity of each HDAC and BDNF band was normalized to the GAPDH reference bands to establish a normalization factor, which was then used to calculate the normalized intensity of the HDAC and BDNF bands in each sample.

### Chromatin immunoprecipitation

Chromatin was prepared using the TruChIP tissue shearing kit (Covaris, Woburn, MA). For each sample, 10 mg of hippocampal tissue was minced into 1 mm3 fragments on dry ice, washed with 1X cold PBS, and centrifuged at 200g for 5min at 4 °C. Pellets were resuspended in Covaris fixing buffer A and fixed in methanol-free formaldehyde (1% final concentration) for 5 mins. After centrifugation, the cell pellet was washed with cold PBS, then flash-frozen in liquid nitrogen and pulverized in a Covaris tissueTUBE. Preparations were lysed in Covaris lysis buffer, and nuclei were sheared for 8 mins on a Covaris M220 instrument. From each sample, 2% of total chromatin was retained, and crosslinks were reversed to provide an input control. The remaining chromatin was split into two equal portions, and ChIP was carried out in ChRIPA buffer after SDS sequestration using either anti-H3K9ac antibody (39137, Active Motif, Carlsbad, CA) or Rabbit IgG isotype control (ab171870, Abcam). Samples were first precleared with anti-rabbit IgG M280 Dynabeads (Thermo Fisher, Waltham, MA) for 1 hour, then incubated overnight with the antibody at 4 °C. The formed complex was captured using fresh Dynabeads for 4 hours at 4 °C. The bead-linked complex was captured by a magnet and washed 4 times with LiCl wash buffer before elution. DNA was liberated from the ChIP’ed samples and input controls by overnight incubation (16 hours) at 65°C, followed by purification using the Minelute kit (Qiagen, Hilden, Germany).

H3K9ac occupancy at the *Bdnf* promoter IV was assessed by quantitative PCR (qPCR) using primers and conditions previously described (Seo et al. 2016; Asp 2018). 2.5 µL of each DNA elution (H3K9ac, IgG, and input control) per sample was used per qPCR reaction. Each sample was amplified in triplicate on a QuantStudio 3 (Applied Biosystems, Foster City, CA) using PowerUp SYBR Green reporter dye. A single melt peak was observed in the qPCR melt curves, indicating the formation of a single product. Standard curves of serially diluted rat DNA were also run for this primer set to evaluate performance, and the standard curve showed an excellent fit (R^2^ = 0.99).

### Data analysis for Westerns and ChIP

Data were analyzed, and graphs were plotted using GraphPad Prism (Version 9). Independent t-tests or a one-way Analysis of Variance (ANOVA) were used to determine statistical significance, indicated by *p < 0.05.

### RNA sequencing (RNA-seq)

Total RNA was isolated using the RNeasy Lipid Tissue Kit (Qiagen, Hilden, Germany). RNA integrity (RIN) was assessed using the RNA 6000 nano kit on a BioAnalyzer (Agilent Technologies, Santa Clara, CA), yielding a mean RIN of 8.8 and a range of 8.4-9.2. Polyadenylated RNA was enriched using the NEBNext® Poly(A) Magnetic Isolation Module (New England Biolabs, Ipswich, MA), and sequencing libraries were prepared from it using the IDT Biosciences RNA library kit (Integrated DNA Technologies, Coralville, IA). Samples were sequenced PE150 using standard Illumina chemistry (Illumina, San Diego, CA). Raw data as FASTQ files were analyzed using FastQC v0.11.9 (quality base calls, CG content distribution, duplicate levels, complexity level) (Andrews 2010). Sequencing adapters were removed using TrimGalore v.0.6.4_dev (Krueger et al. 2023). Raw sequence data are available to download from the Sequence Read Archive at the National Center for Biotechnology Information.

Reads were aligned to the Rn7 rat reference genome using STAR v.2.5.2b (Dobin et al. 2013) and counted on the gene level using featureCounts v2.0.1, using the transcript annotation Rattus_norvegicus.mRatBN7.2.108.chr.gtf. Principal component analysis was applied to RNA-seq data to evaluate the overall similarity and differences among samples. RNA-seq counts were preprocessed and analyzed for differential expression using edgeR v.3.30.0 (Robinson et al. 2010). P-values for differentially expressed genes were corrected using a False Discovery Rate (FDR) threshold of 5% (Benjamini and Hochberg 1995).

### Pathway analysis of RNA-seq findings

To identify common biological themes among the significantly differentially expressed genes from RNA-seq, the top transcripts were subjected to pathway analysis using the online resource at GeneOntology.org (Mi et al. 2017). The organism was specified as *Rattus norvegicus,* and we used Gene Ontology (Gene Ontology Consortium et al. 2023) and PANTHER version 18.0 (PANTHER Overrepresentation Test, Released 20231017) with an FDR correction of 5%. Plots were made using RStudio running R version 4.1.3 or later.

## Results

### Selective increases in hippocampal HDAC proteins in CPF rats

We previously showed increased levels of hippocampal HDAC1 and HDAC5 following exposure to OP compound DFP (Ribeiro et al. 2021; Ribeiro-Davis et al. 2024). We assessed expression of these HDAC proteins in hippocampal homogenates from CPF-exposed and age-matched control rats. In addition to HDAC1 and HDAC5, we also assayed HDAC2 and HDAC3 because of their functional importance in the hippocampus (Guan et al. 2009; McQuown et al. 2011). As shown in **Fig. 2**, significant increases were observed in the levels of HDAC3 and HDAC5 proteins in the CPF rat hippocampus, with densitometric analysis revealing an approximately two-to threefold increase in HDAC expression compared to age-matched control rats (n=5 rats/group, t-test, *p*= 0.01). We observed no significant changes in HDAC1 and HDAC2 levels between CPF and control subjects (data not shown).

**Figure 2.**
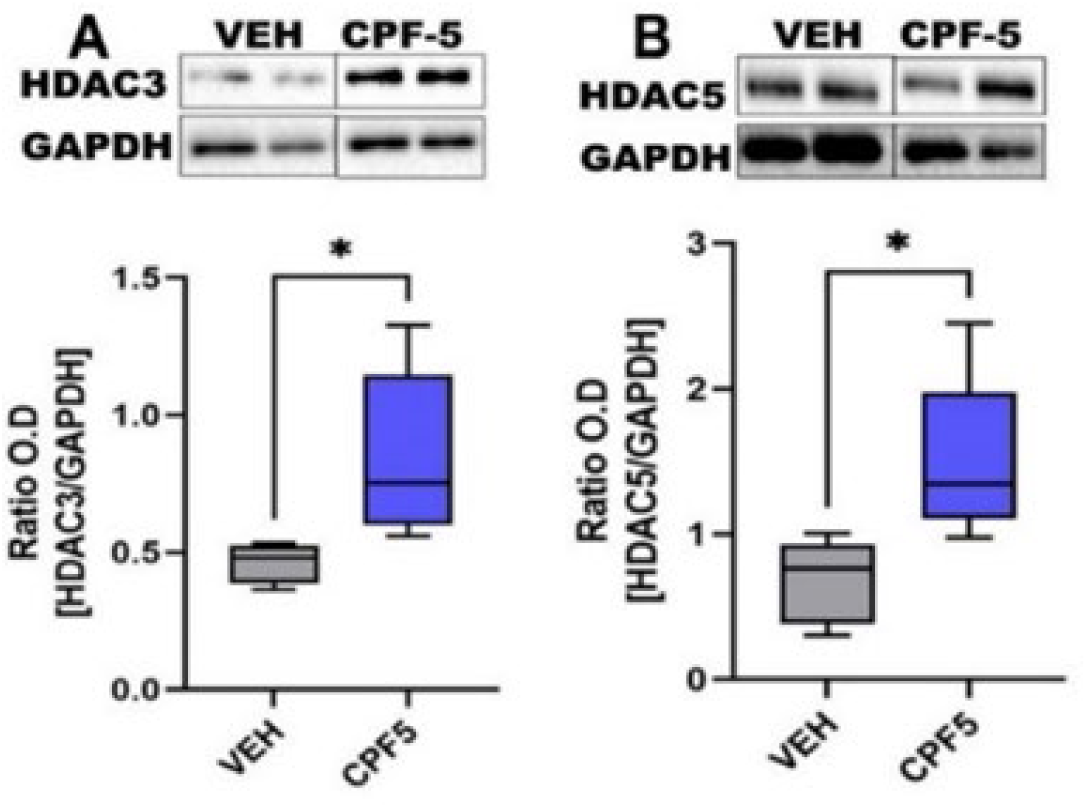
Selective increases in hippocampal histone deacetylase levels following CPF exposure. Western blot analysis showed increases in **(A)** HDAC3 and **(B)** HDAC5 protein levels in CPF-5 rats compared to VEH-treated age-matched controls (n=5 rats/ group, t-test). Representative immunoblots and densitometric analyses are shown.

### Decreased histone acetylation at the *Bdnf* promoter and reduced BDNF expression in CPF rats

HDAC5 has been shown to directly affect histone acetylation levels at the brain-derived neurotrophic factor (*Bdnf*) gene promoter IV (Yadav et al. 2024). We tested if acetylation of histone 3 lysine 9 (H3K9ac) was altered at *Bdnf* promoter IV using ChIP-qPCR. H3K9ac is one of the primary acetylation marks in promoters that is associated with activated gene expression (Tessarz and Kouzarides 2014). As shown in **Fig. 3A**, we observed a significant decrease in H3K9ac levels at the *Bdnf* promoter IV in CPF rats as compared to controls (percent input material 1.00 ± 0.04 (CTL) vs 0.64 ± 0.1(CPF), n = 5 rats/group, t-test, *p* = 0.005).

**Figure 3.**
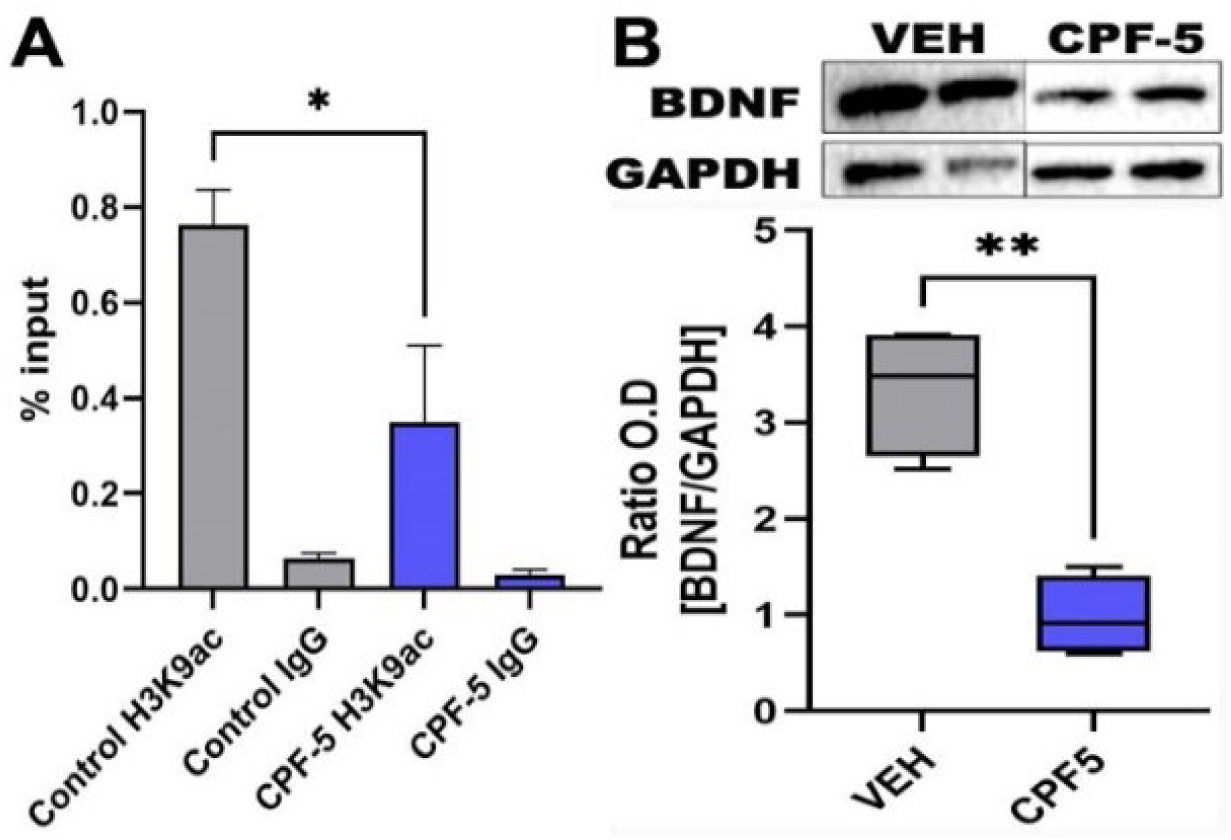
Reduced H3K9 acetylation at the *Bdnf* promoter IV and BDNF expression following CPF exposure. **(A)** ChIP-qPCR showed a reduction in H3K9ac occupancy at *Bdnf* promoter IV in CPF-5 rats compared to VEH. **(B)** Western blot analysis revealed a significant decrease in hippocampal BDNF protein levels in CPF-5 compared to VEH (n=5 rats/ group, t-test).

To determine if this chromatin alteration was associated with reduced BDNF expression, we assayed BDNF protein using Western blot analysis. This revealed significant downregulation of BDNF protein expression in hippocampal homogenates from CPF-treated rats compared with age-matched control rats. Densitometric analysis indicated that BDNF levels in CPF rats were approximately 40% of the levels seen in age-matched controls (n=5 rats/ group, t-test, *p* = 0.01, **Fig. 3B**).

### RNA-seq analysis of CPF-induced gene dysregulation in the hippocampus

The observed changes in HDAC enzymes and H3K9ac levels at the *Bdnf* promoter suggested that CPF could be dysregulating other genes as well. Therefore, to test this, we conducted exploratory RNA-seq analysis of hippocampal RNA extracted from CPF-exposed rats and controls (n=4 per group). Samples were subjected to Illumina PE150 sequencing and, following alignment to the rat reference genome, we obtained an average of 40.5M uniquely aligned read pairs (range 29.6M-48.5M) per sample, with an average unique alignment rate of 87.9% (range 85.4-88.5%).

Overall, in our RNA-seq analysis, we identified 100 significantly differentially expressed genes (DEGs) between CPF rats and controls after multiple testing correction (FDR < 0.05). Of these 100 DEGs, 40 were upregulated in CPF rats relative to controls, and 60 were downregulated. Of the 100 DEGs, 93 showed a |log_2_ fold change| > 1, indicating at least a twofold increase or reduction in expression relative to controls. A volcano plot of the results is shown in **Fig. 4**. Notable among the significant DEGs were those implicated in synaptic plasticity, neuronal development, and trophic signaling, such as insulin-like growth factor 2 (*Igf2*), fibroblast growth factor 14 (*Fgf14*), and slit guidance ligand 2 (*Slit2*). All three genes were significantly downregulated in the CPF group (FDR<0.05), with *Igf2* showing a 6.1-fold reduction, or 16.5% of the expression of the control group (log_2_FC = -2.602), *Slit2* showing a 5.6-fold reduction, or 18% of the expression of the control group (log_2_FC = -2.476), and *Fgf14* showing a 3.3-fold reduction, or 30.3% of the expression of the control group (log_2_FC = -1.721).

**Figure 4.**
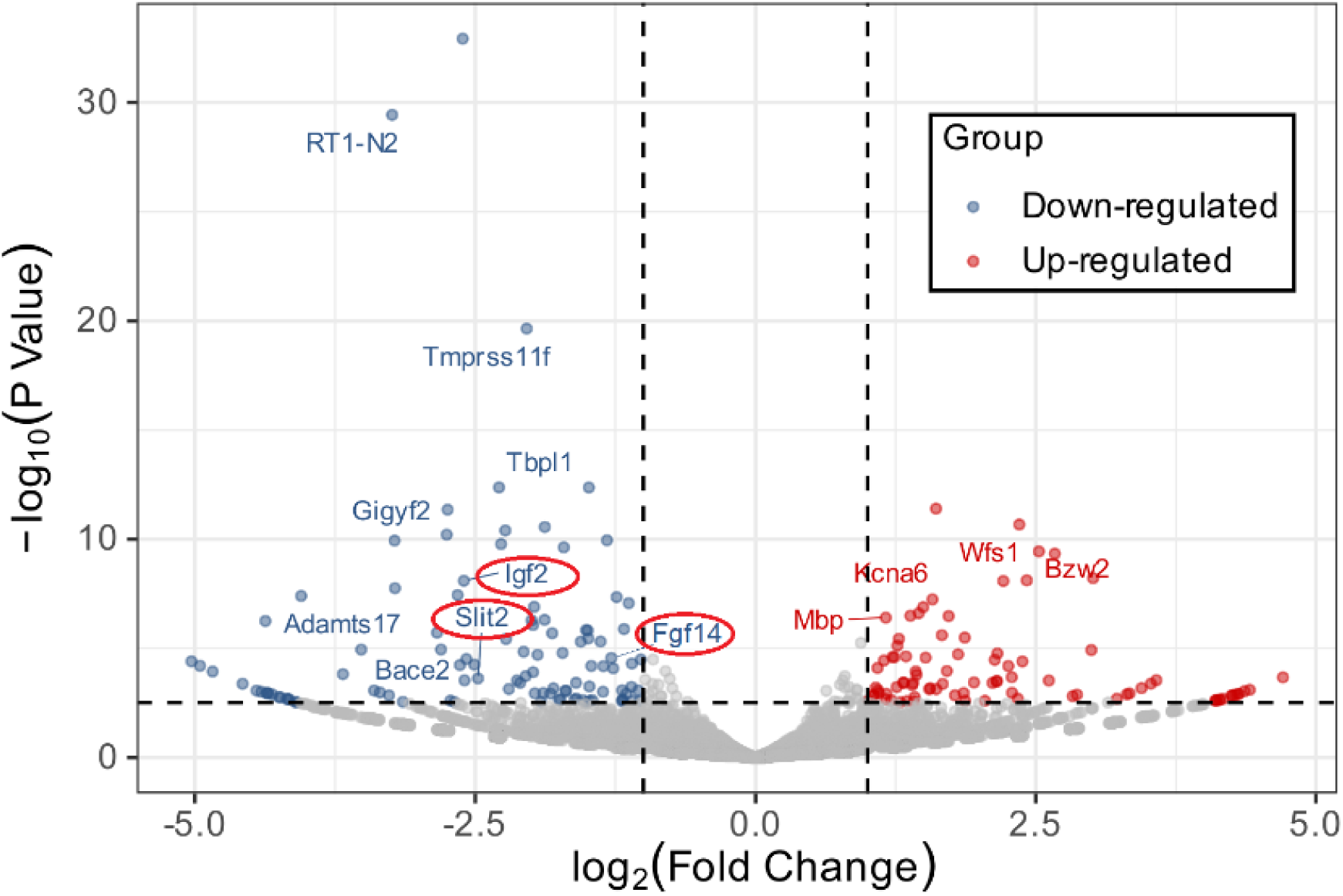
Chronic hippocampal transcriptomic dysregulation following CPF exposure. RNA-seq was performed on hippocampal RNA obtained from CPF-5 and VEH rats. We obtained ∼30M PE150 clusters per sample (n=4/ group). A volcano plot of differentially expressed genes is shown. Vertical dotted lines indicate a 2-fold up/down change in expression, i.e., |Log_2_(Fold Change) | ≥ 1; the horizontal dotted line indicates the FDR < 0.05 significance threshold. Up-regulated genes are shown in red, while down-regulated genes are shown in blue. Down-regulated genes with well-characterized involvement in trophic support, synaptic function, and neuronal development have been highlighted.

### Gene Ontology analysis of CPF differentially expressed genes

To test for common biological themes among the DEGs, we looked for their enrichment in specific gene ontologies (GOs). **Fig. 5** shows the GO terms that were significant at FDR < 0.05. Overall, the most significant GO was “negative regulation of neuron projection development” (GO:0010977, *p* = 6.11×10^-05^). The GO showing the most DEGs relative to the overall set size was “regulation of neurogenesis” with 13 genes out of the full set size of 107 detected in our data (GO:0050767, *p* = 8.96×10-05). Overall, several significant GO terms were related to the themes of nervous system development, myelination/axons, and neurogenesis.

**Figure 5.**
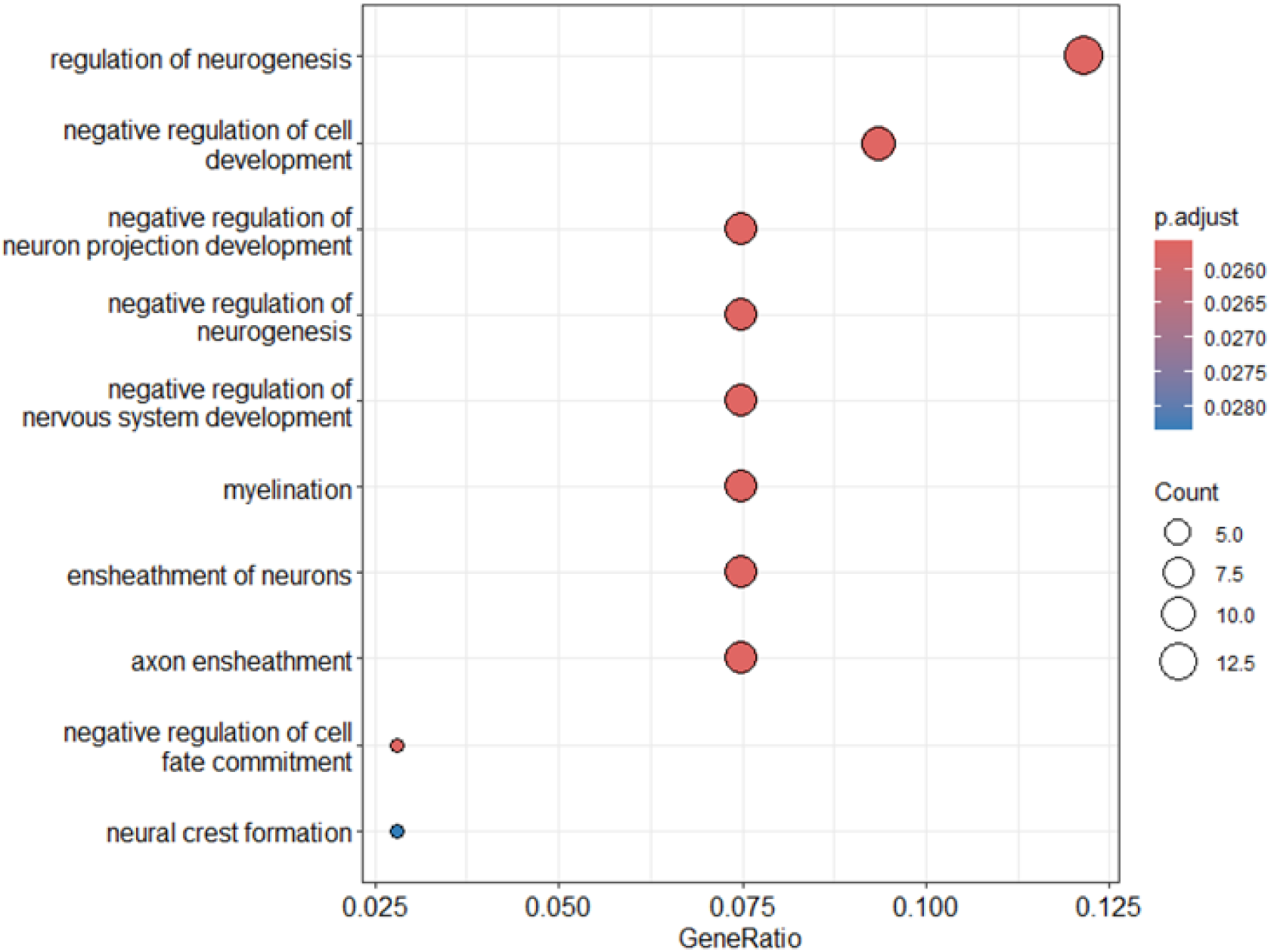
Gene Ontology analysis of differentially expressed genes associated with CPF exposure. We detected 100 differentially expressed genes with FDR < 0.05. Pathway analysis was carried out on these 100 genes, and the top-enriched Gene Ontologies are shown in the bubble plot. The gene enrichment ratio is shown on the x-axis, while the number of genes per ontology is indicated by bubble size. Significance after adjusting for FDR <0.05 (p.adjust) is color-coded, with red indicating greater significance, as shown in the legend. All ontologies shown surpassed the FDR<0.05 threshold.

## Discussion

CPF is now listed as a persistent organic pollutant under the Stockholm Convention (Stockholm Convention on Persistent Organic Pollutants 2025). Human biomonitoring studies demonstrate widespread exposure to CPF, with its urinary metabolite TCPy detected in 82% of the US adult population in early national surveys and in 98.8% of the nationally representative US population in the recent NHANES survey (Hill et al. 1995; Trunnelle et al. 2014; Werthmann et al. 2024). Given the well-established mental health risks of OP exposure (Ross et al. 2013; Beard et al. 2014; Harrison and Mackenzie Ross 2016), coupled with emerging evidence linking exposures to neurodegenerative diseases (Hayden et al. 2010; Voorhees et al. 2019; Hasan et al. 2025), it is vital to look beyond CPF regulatory policy and study the toxicogenomics of sustained low-dose CPF exposures to understand the molecular mechanisms leading to OP-induced neurological morbidities.

In this study, we investigated chronic molecular alterations that persist after repeated CPF exposure. Using hippocampal tissue collected three months after occupational-like CPF exposure from rats that exhibited a depressive-like phenotype, we identified selective HDAC dysregulation, reduced histone acetylation at the *Bdnf* promoter IV accompanied by sustained BDNF downregulation, and broader transcriptional alterations involving neuronal plasticity-related pathways. The persistence of molecular and behavioral abnormalities despite recovery of cholinesterase activity supports the involvement of non-cholinergic mechanisms in the long-term consequences of CPF exposure.

Previous research has identified diverse molecular targets of CPF, including calcium dysregulation, mitochondrial toxicity, oxidative stress, neuroinflammation, and cytoskeletal protein impairment, as potential non-cholinergic mechanisms underlying long-term OP neurotoxicity (Pope 1999; Terry 2012). In addition to these cellular mechanisms, disruption of neurotrophic signaling has emerged as a critical non-cholinergic mechanism underlying the persistent neurobehavioral deficits following OP exposures. Thus, using a CPF exposure regimen similar to that in our study but examining the tissue one day after exposure, Lee et al. (2016) reported alterations in hippocampal gene expression of neuropeptides and neurotrophins. Additionally, using similar CPF doses but with only a 5-day exposure duration, Speed et al. (2012) reported delayed hippocampal synaptic abnormalities in mice. Furthermore, in a rat model of Gulf War Illness that used repeated, low-dose exposure to the OP-based nerve agent surrogate DFP (diisopropylfluorophosphate), we reported mood and memory impairment alongside sustained BDNF downregulation (Ribeiro et al. 2021; Ribeiro-Davis et al. 2024). Given the critical role of BDNF in neuroplasticity and mood regulation, understanding the impact of CPF exposure on its regulation can help elucidate how OPs contribute to depressive pathophysiology.

BDNF is an established mediator in the pathophysiology of depression and antidepressant response. There is evidence for reduced BDNF levels in major depressive disorder (Castrén and Monteggia 2021), and this phenotype is replicated in laboratory models of depression (Krishnan and Nestler 2011), including OP-induced depression (Ribeiro et al. 2020; Ribeiro et al. 2021). In agreement with these findings, our study demonstrated that a significant reduction in hippocampal BDNF expression accompanied depression following occupational-like CPF exposures. Given that BDNF plays a vital role in several synaptic plasticity processes, including spine maturation, neurogenesis, and modulation of long-term potentiation (Karpova 2014; Kowiański et al. 2018), this downregulation represents a critical vulnerability in neuronal plasticity, affecting the structural and functional integrity of mood-regulating circuits in the hippocampus.

We next investigated mechanisms for chronic reductions in BDNF levels. For this, we focused on the epigenetic consequences of OP exposures and their effects on gene expression. Our results here demonstrate that occupational-like CPF exposures that produced a depressive phenotype also exhibited changes to epigenetic histone signaling. We noted significant increases in hippocampal HDAC proteins, including HDAC3 and HDAC5, but not HDAC1 and HDAC2. Interestingly, in another OP rat model using the higher potency compound DFP, we observed increases in hippocampal HDAC1 and HDAC5, but no significant changes in HDAC2 and HDAC3 (Ribeiro-Davis et al. 2024). Thus, increased hippocampal HDAC5 expression was common to both OP exposure paradigms, whereas alterations in HDAC1 and HDAC3 differed between the models. These findings suggest that OP-induced epigenetic dysregulation may involve both shared and exposure-specific HDAC responses, with HDAC5 emerging as a potential common component.

The function of HDACs is to deacetylate histones, thereby condensing chromatin and repressing gene transcription. HDAC5 has been shown to affect histone acetylation levels at the *Bdnf* gene (Tsankova et al. 2006) and it binds directly at *Bdnf* promoter IV (Yadav et al. 2024). Therefore, we next investigated if CPF-induced changes to histone acetylation affected *Bdnf* transcription. The *Bdnf* gene contains several differentially regulated promoters, of which promoter IV is the best-characterized regulatory element (Lyons and West 2011). Lack of *Bdnf* expression through promoter IV results in depressive phenotypes in mice (Tsankova et al. 2006; Sakata et al. 2010), while epigenetic modifications at promoter IV are sensitive to stressors and can affect the regulation of genes involved in monoamine neurotransmission (Boersma et al. 2014; Sakata and Duke 2014). Alterations in BDNF levels due to epigenetic modifications were reported in rodent models of GWI (Kimono et al. 2020; Ribeiro et al. 2021). Our ChIP analysis indicated a significant reduction in H3K27ac at the *Bdnf* promoter site on exon IV. Acetylated histones are associated with a relaxed chromatin conformation that allows the cellular transcriptional machinery to access DNA (Tessarz and Kouzarides 2014). Therefore, a reduction in H3K27ac, as observed here, indicates more condensed chromatin and reduced gene expression. Thus, histone deacetylation could explain the reductions in BDNF protein levels seen in this study. Taken together, the increased HDAC5 expression and reduced BDNF levels common to both OP exposure models suggest that HDAC5-associated epigenetic regulation of *Bdnf* represents a shared molecular response across distinct OP exposure paradigms linked to long-term depressive outcomes.

The observation of an altered epigenetic state at *Bdnf* and reduced expression of its encoded protein raised the possibility that CPF may also produce more widespread genomic dysregulation and affect other genes. Therefore, we conducted exploratory RNA sequencing of hippocampal RNAs from CPF-exposed and control rats. This RNA sequencing revealed numerous differentially expressed genes, including downregulation of genes involved in trophic support, synaptic function, and neuronal development. Among these were insulin-like growth factor 2 (*Igf2*), fibroblast growth factor 14 (*Fgf14*), and slit guidance ligand 2 (*Slit2*). Neurotrophic factors such as BDNF, IGF2, and FGF14, and axon guidance molecules such as SLIT2, are critical for dendritic spine maturation and healthy synaptic networks, and reduced levels of these factors are reported in depression (Luo et al. 2015; Alshammari et al. 2016; Li et al. 2017). Consistent with these gene-level changes, our pathway analysis revealed that the differentially expressed genes clustered in biological processes related to nervous system development, myelination, and, notably, the regulation of neurogenesis. These findings are particularly relevant in the context of the sustained reduction in hippocampal BDNF observed in the present study, given the established role of BDNF signaling in neuronal plasticity, neurogenesis, and the maintenance and remodeling of neuronal processes. Indeed, as discussed earlier, we reported that exposure to other OP compounds similarly decreases BDNF levels and reduces synaptic spine density in rodents (Ribeiro-Davis et al. 2024).

There are limitations to this study. While our results support a role for HDAC3/ HDAC5 in *Bdnf* dysregulation, their direct involvement in the loss of H3K9ac at *Bdnf* promoter IV remains to be determined in this model. Studies using ChIP-seq are planned to test whether CPF exposure increases the recruitment of these HDACs to *the Bdnf* promoter *IV* region. Second, we did not detect reduced *Bdnf* transcript levels in our RNA-seq, despite observing a reduction in BDNF protein levels. This is most likely due to our RNA-seq being unable to resolve the large number of alternatively spliced transcripts in *Bdnf* (Maynard et al. 2017). Future studies should employ technologies to assay specific *Bdnf* isoforms. Third, studies for assessing functional synaptic plasticity in CPF-induced depression and upon manipulation of HDACs are needed to provide a molecular framework linking occupational-like CPF exposures to the development and persistence of long-term depression. Fourth, we need to demonstrate alterations in neurogenesis or myelination as identified by our RNA-seq and Pathway Analysis. Finally, the present study utilized only male rats. Sex-dependent differences in OP neurotoxicity have been reported (Comfort and Re 2017). Additional studies are needed to examine whether these CPF-induced long-term epigenetic effects also occur in females.

In conclusion, our experimental studies put forth a model for the development of chronic depression following occupational-like exposures to the OP pesticide CPF during adulthood. We posit that CPF exposures induce epigenetic changes associated with persistent BDNF dysregulation, as well as transcriptional alterations in gene networks involved in neurogenesis, neuronal development, and myelin- and axon-associated processes. Such chronic alterations in neurotrophic signaling could underlie synaptic plasticity abnormalities, thereby contributing to the expression of chronic depression following CPF exposure.

## Acknowledgements

The Genomics Core facility at Virginia Commonwealth University provided access to equipment for sequencing library quality control. High-Performance Computing resources provided by the High-Performance Research Computing (HPRC) core facility at Virginia Commonwealth University (https://hprc.vcu.edu) were used for conducting the research reported in this work. Illumina sequencing was conducted by Azenta/GeneWiz of South Plainfield, New Jersey.

## Author Contributions

AR-D: Formal analysis, Investigation.

EH: Investigation.

FJ: Investigation.

LD: Conceptualization, Methodology, Formal analysis, Writing-original draft, review & editing, Supervision, Project administration, Funding acquisition.

JM: Formal analysis, Writing-review & editing, Supervision, Project administration.

MD: Formal analysis.

